# Promoting synthetic symbiosis under environmental disturbances

**DOI:** 10.1101/395426

**Authors:** Jai A. Denton, Chaitanya S. Gokhale

**Affiliations:** Genomics & Regulatory Systems Unit, Okinawa Institute of Science & Technology, 5 Onna-son, Japan; Research Group for Theoretical Models of Eco-evolutionary Dynamics, Department of Evolutionary Theory, Max Planck Institute for Evolutionary Biology, August Thienemann Str. 2, 24306, Plön, Germany

## Abstract

By virtue of complex interactions, the behaviour of mutualistic systems is difficult to study and nearly impossible to predict. We have developed a theoretical model of a modifiable experimental yeast system that is amenable to exploring self-organised cooperation while considering the production and use of specific metabolites. Leveraging the simplicity of an artificial yeast system, a simple model of mutualism, we develop and test the assumptions and stability of this theoretical model. We examine how one-off, recurring and permanent changes to an ecological niche affect a cooperative interaction and identify an ecological “Goldilocks zone” in which the mutualism can survive. Moreover, we explore how a factor like the cost of mutualism – the cellular burden of cooperating – influences the stability of mutualism and how environmental changes shape this stability. Our results highlight the fragility of mutualisms and suggest the use of synthetic biology to stave off an ecological collapse.

## Introduction

Life on earth comprises a hierarchy of units of selection. From societies to genes, we find the same patterns of organisation at each of these levels [1]. However, selection is not level-specific. While changes may occur at low levels, such as a single nucleotide or amino acid, selection necessarily operates at a much higher level, such as the organism. Competition between entities at a specific level of organisation can spell disaster for the higher level. A clear example of this breakdown of control being cancer [1].

Mutualistic interactions, a specialised type of cooperation where replicating components benefit each other, therefore, can be targeted by selection at a higher level [2]. Due to conflicts of interest between entities at varying levels of selection, the origin of mutualism and subsequent selection lack a clear evolutionary explanation – a complete field of research in itself [3]. Even though it is hard to explain how mutualisms emerge, we see many examples of them. On a global scale, prominent examples such as coral-*Symbiodinium* symbioses or plant-rhizobia interactions are well known [4, 5, 6]. Many such mutualisms have evolved over millions of years, but if mutualisms are fragile and susceptible to collapse, as hypothesised, then how do they survive for aeons in constantly changing environments?

Mutualistic interactions can emerge in numerous ways [3]. Here we focus on how they survive, because, regardless of how they originate, mutualisms constantly face different threats. A common challenge for mutualistic communities is exploitation by cheater strains that benefit from mutualistic interactions, but fail to contribute.

This problem of parasitic elements, first noted by Maynard Smith [7], has been extensively studied. Postulations about compensatory mechanisms that could avoid parasitic exploitation, range from conceptual [8], to mechanistic arguments [9]. Mutualism could also suffer from insufficient ecological support. That is, if population densities of mutualistic partners are inadequate, then mutualisms are hard to sustain. This problem highlights the necessity of a physical structure akin to a “warm little pond” for concentrating the initial mutualists [10]. Without a physical structure or substrate to provide overlapping niches, it would be hard to kick-start necessary mutualistic reactions [11]. However, what happens to the stability of a mutualistic system when a threat to mutualism does not involve a parasitic element, but an unsupportive environment for the system itself? Understanding how cooperative interactions survive in changing environments that continually alter the basis of mutualistic interactions is then worthy of investigation.

The selective advantage derived from cooperation may be transitory in the face of environmental changes; therefore, in most cases the dynamics of such systems remain in constant states of flux. Interspecific interactions are essential in determining community stability [12]. Depending on resource availability, interactions between mutualists can change to facultative mutualism, to competition, and even parasitism [13]. In a synthetic system, interactions are fixed; therefore, environmental influences on the system can be determined. However, ecological networks are inherently complex [14]. While a complete understanding of a network provides us with general properties of the interactions, it is often impossible to pin down the principles that form the building blocks. As such, it is easier to study tightly controlled systems with a defined number of interactions. Numerous systems adapted from nature allow us to study evolutionary and population dynamics.

Via genetic manipulation, synthetic, cross-feeding, cooperation can be engineered within microbial communities [15, 16, 17, 13, 18]. Complex population dynamics in response to temporal, spatial, and environmental factors can thus be dissected by fine-tuning and manipulating these synthetic systems [19, 18]. We use *Saccharomyces cerevisiae* synthetic mutualism systems that rely on cross-feeding of amino acids between two strains [15].

The system uses feedback resistance (fbr) mutations in adenine and lysine biosynthesis pathways that results in overproduction of the corresponding amino acid. The strains used are referred to here as *LYS↑* and 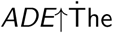 dynamics of this cross-feeding system is comparableto a simple, but powerful theoretical model of self-organisation – a hypercycle – implicated at the origin of life as well as the formation of complex communities [20, 21, 11, 22, 18]. Here we develop a simple, but powerful and easily extendable phenomenological model. We then use the yeast system to validate the model and finally explore beyond the synthetic system to understand stability and cost of cooperative interactions in changing environments. While it is clear that a change in the environment affects the interaction pattern, our aim is to dissect abiotic factors to under how exactly this occurs. With this aim in mind we begin with our theoretical and experimental model.

## Model & Results

Our model begins with the growth of two yeast strains in an environment that lacks free adenine and lysine as nutrient sources. The two strains are *LYS↑* and *ADE↑* whose densities are denoted by *x*_*L*_ and *x*_*A*_. The strain *LYS↑* is deficient in adenine (density *c*_*A*_) while it overproduces lysine (density *c*_*L*_). The production and requirement for adenine and lysine is reversed for 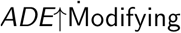 odifying the logistic growth equation gives the dynamical equations for the growth of the two strains:

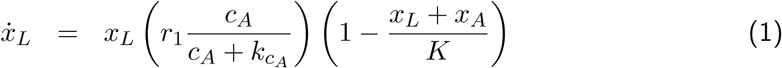

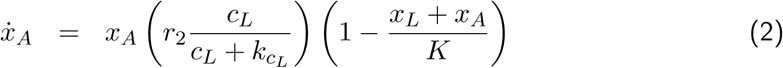

The two strains grow if the required metabolites are present. The strains compete for a limited amount of space, given by *K*. Amino acid concentrations, together with Monod-type saturation kinetics, control the growth of the strains. Our model is mechanistic regarding individual amino acid dynamics. Explicitly including metabolite concentrations is crucial, as pairwise Lotka-Volterra models may not always provide a realistic qualitative picture of the dynamics [14]. Amino acids are a consumable resource. As they are produced constitutively by one of the strains, the other strain uses them immediately. The dynamics of metabolite densities in the culture hence can be captured by:

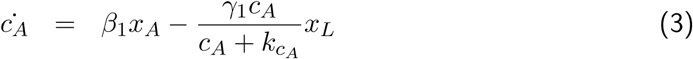

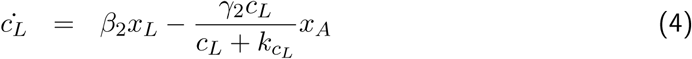

The rate at which each metabolite *i* can increase in the yeast culture is given by *β*_*i*_. This rate determines the amount of metabolite shared by the corresponding overproducing strain. We assume that use of a metabolite at rate *γ*_*i*_ by strains *x*_*i*_ also involves formation of an intermediate; thus being subject to Michaelis-Menten kinetic parameters. The simple dynamics of such a system are depicted in Fig. 1.

**Figure 1:**
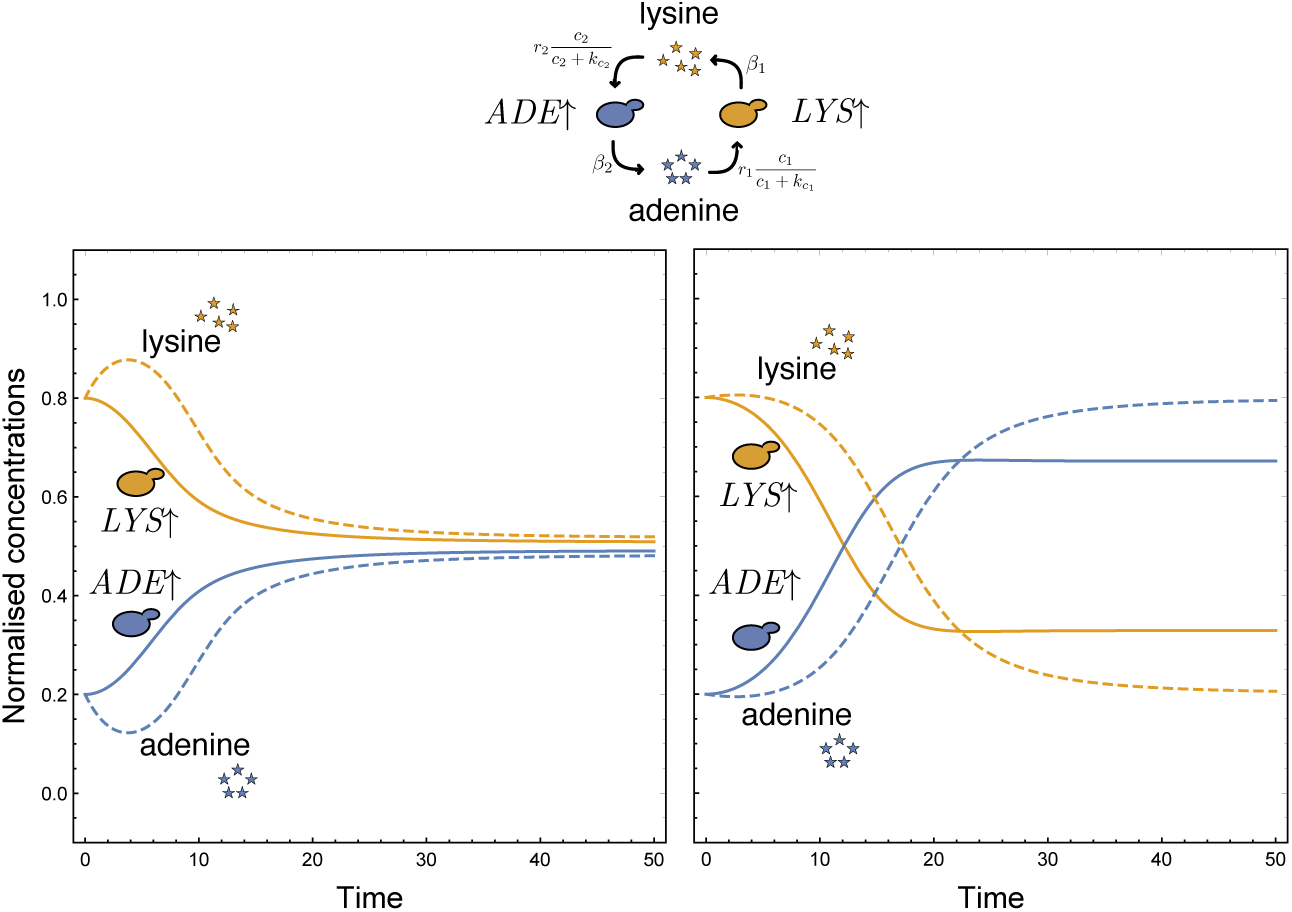
Unsupplemented dynamics (left) the neutral expectation and (right) the expectation informed by experimental growth rates. We plot the fraction of *LYS↑* (*x*_*L*_*/*(*x*_*L*_ + *x*_*A*_)), *ADE↑* (*x*_*A*_*/*(*x*_*L*_ + *x*_*A*_)) and the relative fractions of metabolites adenine (*c*_*A*_*/*(*c*_*A*_ + *c*_*L*_)) and lysine (*c*_*L*_*/*(*c*_*A*_ + *c*_*L*_)) in a continuous culture. As an initial condition there is no free adenine or lysine in the culture, but nonzero populations of the strains generate the free adenine and lysine in the same relative frequencies. The relative initial fractions for the strains are *LYS↑* = 0.8 and *ADE↑* = 0.2 and the rate at which the two strains share the two metabolites is set to *β*_1_ = *β*_2_ = 0.04. We set the carrying capacity of the culture vessel to *K* = 5. **Left:** The uptake rate of the metabolites as well as the degradation rate are exactly the same (*r*_1_ = *r*_2_ = 5 and *γ*_1_ = *γ*_2_ = 2). Michaelis constant for both metabolites to 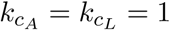. Under these symmetric conditions, normalised densities of the strains, as well as that when the two metabolites reach an equilibrium where normalised concentrations are the same. **Right:** Clearly the assumption of symmetric rates is a simplification. Informed by experiments, we estimate *r*_1_ = 11.4906, *r*_2_ = 29.3955 and the Michaelis constants 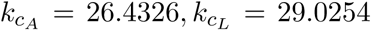. This asymmetry in uptake rates is reflected in the resulting unequal equilibrium.

Adding the required supplement, we calculate the growth rate of the two strains for different levels of supplementation. The resulting growth rate curves are used to parameterise the model. The growth parameters are *r*_1_ = 11.4906 and *r*_2_ = 29.3955 and the Michaelis constants 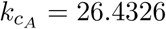 and 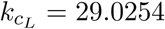. We assume the Michaelis constant estimated for the uptake rate for adenine and lysine to be the same as the rate at which they degrade from the pool. Starting at different initial conditions of the *ADE↑* to *LYS↑* ratio, the final equilibrium values are close to 0.6, corroborated by experiments Fig. 2.

**Figure 2:**
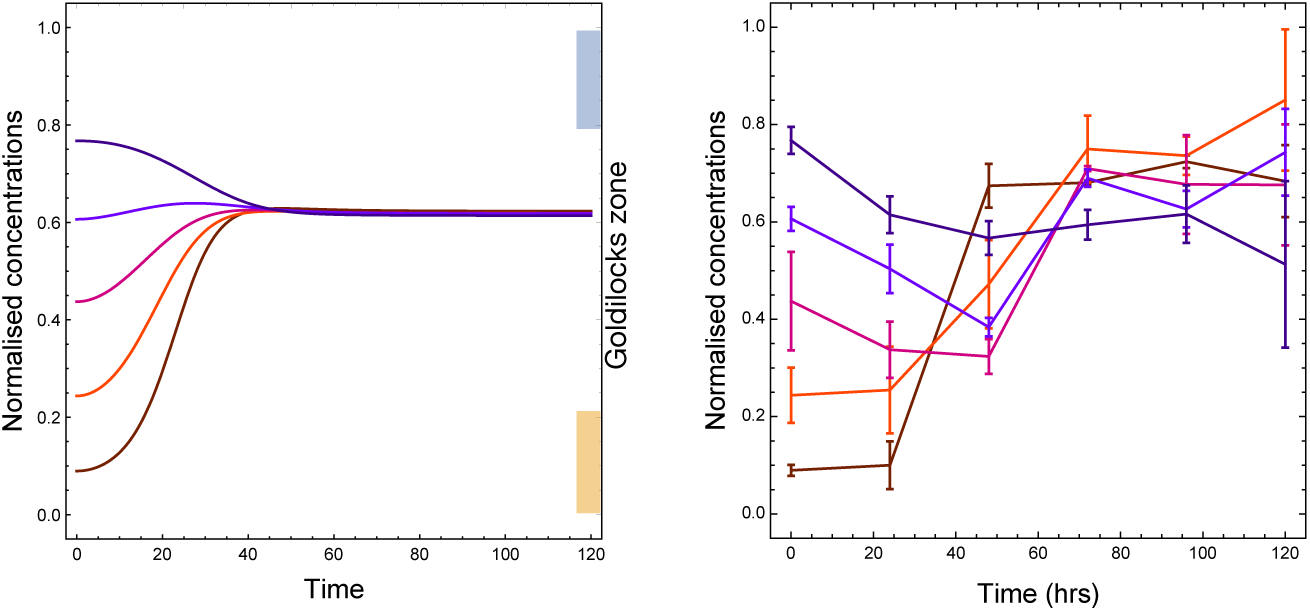
Theoretical and experimental model dynamics of with different initial ADE*↑* concentrations. Using the theoretical model, in the left panel we predict the dynamics of the normalised fraction of *ADE↑* starting at various initial ratios with the parameters set informed by data. The initial concentrations of the amino acids *c*_*A*_ and *c*_*L*_ are set to 0.0. Using the experimental model, in the right panel we plot the the normalised concentration of the yeast strain *ADE↑* relative to the *LYS↑*. Starting at different initial ratios we track dynamics over a 140-hour period in synthetic complete media (without adenine or lysine). Each point is the mean of three replicates and error bars represent a single standard deviation.

## Eco-evolutionary dynamics under environmental disturbance

The mutualistic system rests on the interdependence of the two strains of yeast. If we undermine this dependency, it will affect the stability of the mutualistic interaction. Typically the system can be undermined by the introduction of a parasite [18]. We choose to change the environment itself without altering the genotypes of the interacting strains.

If the environment has one of the two required metabolites, then the strain using that metabolite will not depend on the corresponding strain for survival. A significant amount of supplementation can drown out the signal of the other strain. In all, we visualise three different supplementation scenarios that could be used to test the resilience of this mutualistic system. Specifically, initial supplementation – adenine or lysine added at the beginning of culturing, continuous supplementation – the two metabolites added steadily throughout culturing, and intermittent supplementation – the metabolites added at regular time intervals. Depending on the timescale, a single niche can experience all three scenarios, but we analyse the three scenarios in succession.

## Initial supplementation

The initial supplementation regime is a standard batch culture. With nutrients provided at the start, the culture continues to grow until one or more nutrients become limiting. We supplemented our experimental cultures with either 0.1, 1, or 10 µg/mL adenine and measured the effect on strain ratio at regular time intervals (Materials & Methods, Fig. 3 (top)). The supplementation favoured the *LYS↑* strain, resulting in a lower prevalence of *ADE↑* when compared to the unsupplemented condition. The mean strain ratio of *ADE↑* to *LYS↑* after 120 hours growth of all starting ratios for 0.1, 1, or 10 µg mL^−1^ adenine supplementation is 0.5316, 0.5685, 0.0556, respectively.

**Figure 3:**
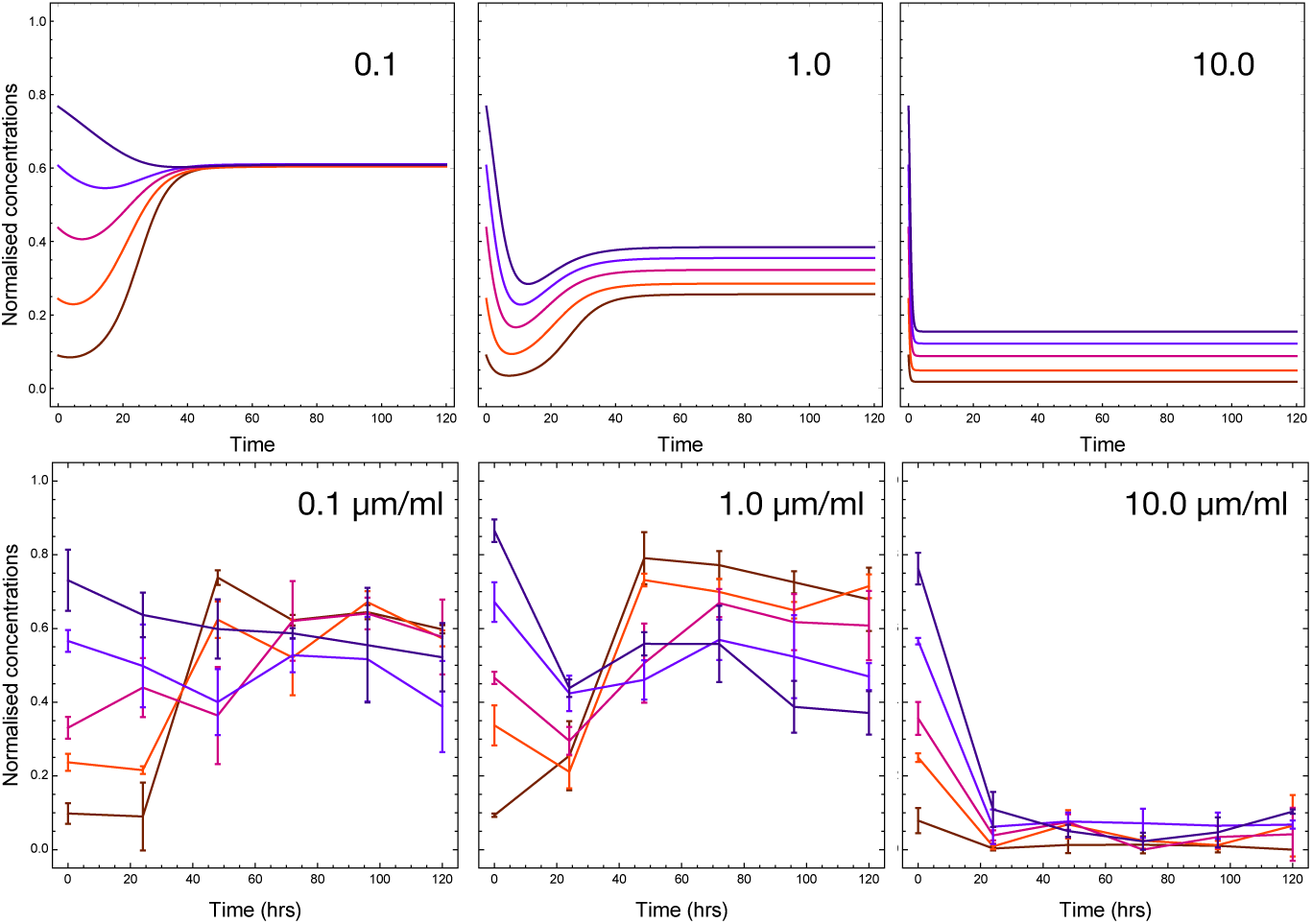
Theoretical and experimental model dynamics with different initial supplementation levels of adenine. We explore the theoretical model as depicted in Fig. 2, now with added initial supplementation. The initial concentrations of the metabolites *c*_*A*_ and *c*_*L*_ are set to 0.1, 1.0, and 10 as denoted in the panels. Using the modified version of In the bottom row we show the normalised concentration of the yeast strain *ADE↑* relative to the *LYS↑* strain, at varying starting ratios over a 120-hour period in synthetic complete media supplemented with adenine to a final concentration of either 0.1, 1.0, or 10 µg/mL. Each point is the mean of three replicates and the error bars represent a single standard deviation. The theoretical model qualitatively predicts experimental model dynamics of mutualism. With increasing supplementation, the dependence of *LYS↑* on *ade*^*fbr*^*lys*^*-*^ is reduced and the system exits the Goldilocks zone.

Using the parameterised theoretical model, we explore the final equilibrium value of *ADE↑* as we change the amount of initially supplied metabolites. The model qualitatively mimics the change in the equilibrium condition with the addition of adenine Fig. 3 (bottom). Experimentally we have tested the addition of only adenine for different initial fractions of *ADE↑* but theoretically, we explore the consequence of adding both adenine and lysine for the *ADE↑* starting at 0.5. The results, summarised in Fig. 4 (left panel), reveal that even a slight asymmetric increase in the amount of initial metabolite present in the environment is enough to destabilise the equilibrium and push it close to extinction. When the equilibrium values come close to one of the two edges of the system, the rarer mutualism runs the risk of going extinct by drift. Close to the edges (0 or 1), stochastic dynamics could lead to the extinction of either strain. If the equilibrium value of either mutualist falls below 0.2, we assume that the mutualist system is at high risk of collapse. Thus we define the region between 0.2 and 0.8 as the “Goldilocks zone” where mutualism can safely exist. The asymmetry of the graph in Fig. 4 reflects the inherently different uptake rates of metabolites by the two strains Fig. 2.

**Figure 4:**
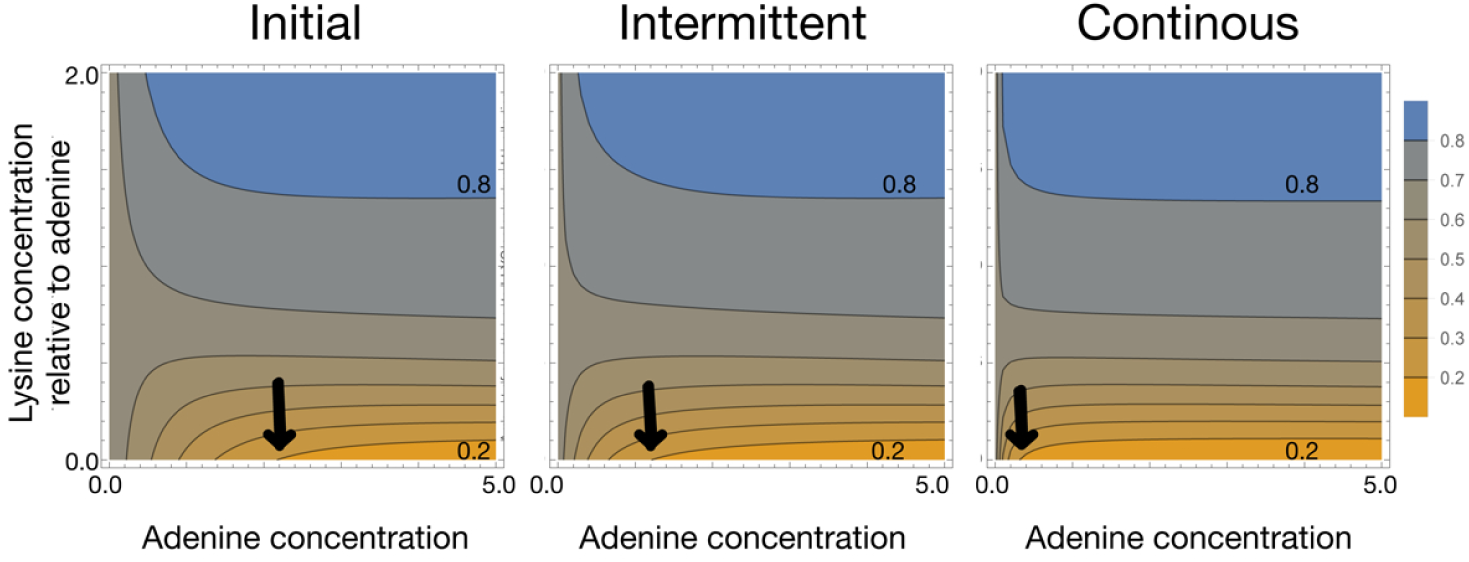
Equilibrium value of ADE*↑* for different supplementation regimes. Initial normalised concentration of *ADE↑* = 0.5. The system is assumed to have equilibrated by 300 time steps. In the plots we show the eventual normalised concentration of *ADE↑* going from all *ADE↑* (1, blue) to all *LYS↑* (0, yellow). The initial concentration of adenine in the system at time 0 is shown on the *x*-axis and the amount of lysing relative to adenine is depicted on the *y*-axis. For *y* = 1 the amount of adenine and lysine is the same. The initial supplementation: amino acids provided only at time point *t* = 0, intermittent supplementation: starting at *t* = 0 and then every 5^*th*^ time step and for continuous: at every time step. Going from Initial to Continuous supplementation we see that the Goldilocks zone (region between the contours 0.2 and 0.8) shrinks as the zone for *LYS↑* (yellow) increases, and so does the *ADE↑* (blue) region.

## Intermittent supplementation

As the initially added supplementation is consumed, it is possible to add metabolites at fixed intervals. The model is now an example of a hybrid dynamic system [23] where concentrations of the metabolites are adjusted at regular intervals (see Materials & Methods).

Intermittent supplementation acts as a bridge between the other two supplementation regimes. If the delay between two supplementations is substantial then starting at time *t* = 0, the scenario is the same as that of initial supplementation (Fig. 4 middle panel would resemble the left panel). If the delay between successive supplementations is minimal, then the concept is similar to continuous supplementation (Fig. 4 middle panel would resemble the right panel). For a legitimate comparison with initial and continuous supplementation, we started the intermittent supplementation with a non-zero level of amino acids which was added at regular intervals.

## Continuous supplementation

Intermittent supplementation, if done at minimal time intervals, represents a continuous supply of amino acids. Continuous supplementation competes with the biosynthetic output of the strains themselves, drowning out their signal and undermining the basis of cooperation between the two strains.

As before, we plot equilibrium values of *ADE↑* for different amounts of continuously added adenine and lysine (relative to adenine concentration) (Fig. 4 right panel). Slight, but continuous addition of lysine immediately breaks the mutualism as the *ADE↑* strain takes over the mixed culture. The Goldilocks zone breaks down almost immediately for continuous supplementation as compared to initial supplementation.

## Intervention measures in the presence of ultimate overproducers

A typical threat to a mutualistic system is the evolution of a cheater, which parasitizes the produced common good [24]. A cheating strain can thrive because it benefits from public goods without having to pay the cost of contributing. But does mutualism always have to be extremely costly? If production of the common good entails little cost, then we could envision mutants that do not require mutualistic interactions, but that contribute to the pool of common goods, like fictitious *ADE↑LYS↑* strains. While not participating in mutualism might be lethal, taking part in a mutualistic interaction does not need to be costly if it involves overproduction of a single metabolite. Although specific, yeast can overproduce a fluorescent protein and only suffer a 1% reduction in cost per copy [25]. Moreover, reliance on external sources for essential metabolites can have a considerable cost as well. Several auxotrophic yeast strains, such as those unable to produce their own lysine or adenine, have up to a 10% reduction in growth, even with environmental supplementation [26, 27].

Typically if a strain overproduces a compound, a cost is associated with it. However, there is potentially also a cost associated with relying on interactions with other organisms. Our mathematical model is also extensible to include a fictitious strain *ADE↑LYS↑* that overproduces both amino acids and does not require any supplementation. We assume that any cellular cost incurred will not matter unless it affects the growth rate. In the simplest case, the growth rate would be largely independent of the environment since the strain can satisfy its requirements.

Freeing itself from any environmental dependence on metabolites, our ultimate overproducer is defined as having a variable flat growth rate. Compared to the growth rates extrapolated from the experimental data we can envision a number of different scenarios as in Figure 6 (top). The variable growth rates chosen for the overproducing strain reflect a range of values within those derived from the other strain experimental data. A clear prediction for which of the three strains is dominant is derived from our updated model and closely reflects the underlying growth rates. Even with a comparatively high cost, the *ADE↑LYS↑* strain can quickly become the dominate strain, despite supporting both the *ADE↑* and *LYS↑* strains. If we are interested in ensuring mutualism, then we must intervene, but the question is when and to what degree?

**Figure 5:**
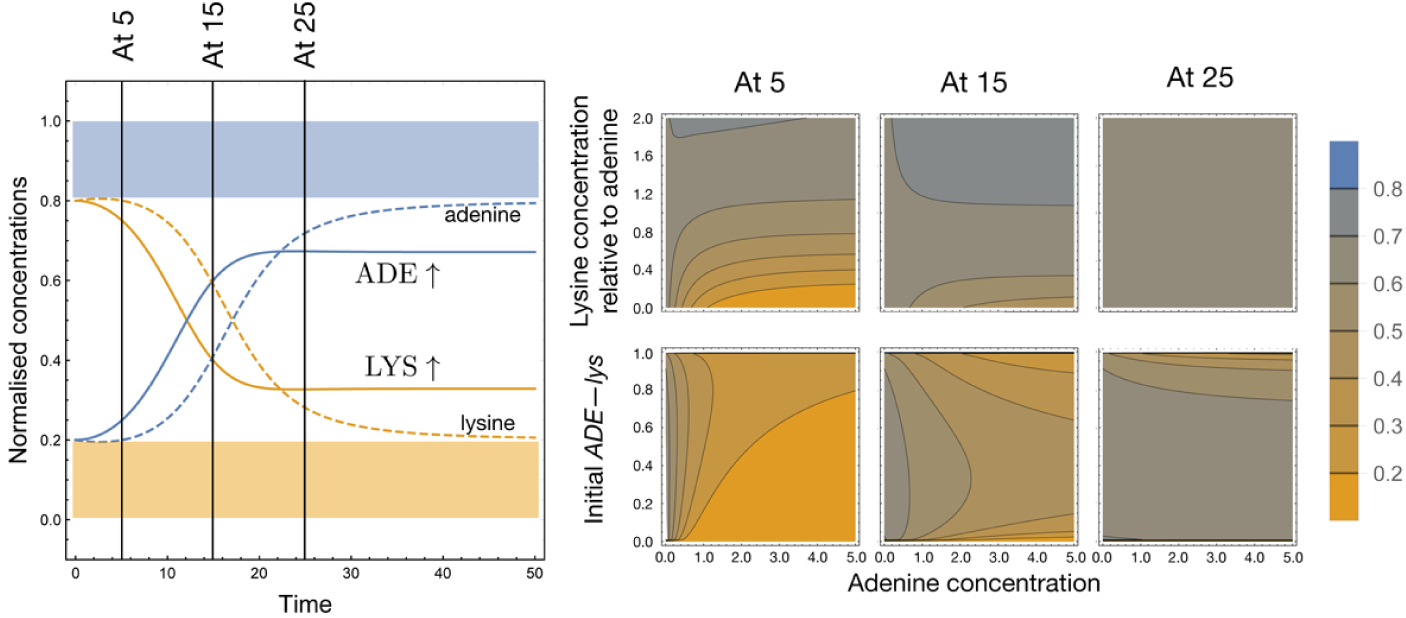
Disrupting transient mutualism with intermittent supplementation. Intermittent supplementation (Figure 4) started with the same initial supplementation as the initial supplementation regime, so as to be comparable. If we start with no metabolites in the culture and add them at fixed intervals, then it might be possible to extend the range of the Goldilocks zone, even under supplementation. The exact timing of supplementation provided is crucial in determining the eventual equilibrium. If the pattern of supplementation starts early in the existence of *ADE↑* then the resulting equilibrium frequency can be drastically affected. By tuning the timing and dose of supplementation, we can maximise the probability of maintaining the system within the Goldilocks zone (as we move from supplementing every 5 to 15 to 25 time step). If intermittent supplementation starts in the latter phase of the transient where the equilibrium value is already reached then the co-existence seems robust. This will also depend on the initial concentration of *ADE↑*. Thus, while the top row is calculated from an initial normalised concentration of *ADE↑*= 0.2, the bottom row explores all initial conditions. Equilibrium values are all calculated at the same time point, i.e., the number of cycles are adjusted so that all integrations run until time point *cycle length × number of cycles* = 300.

**Figure 6:**
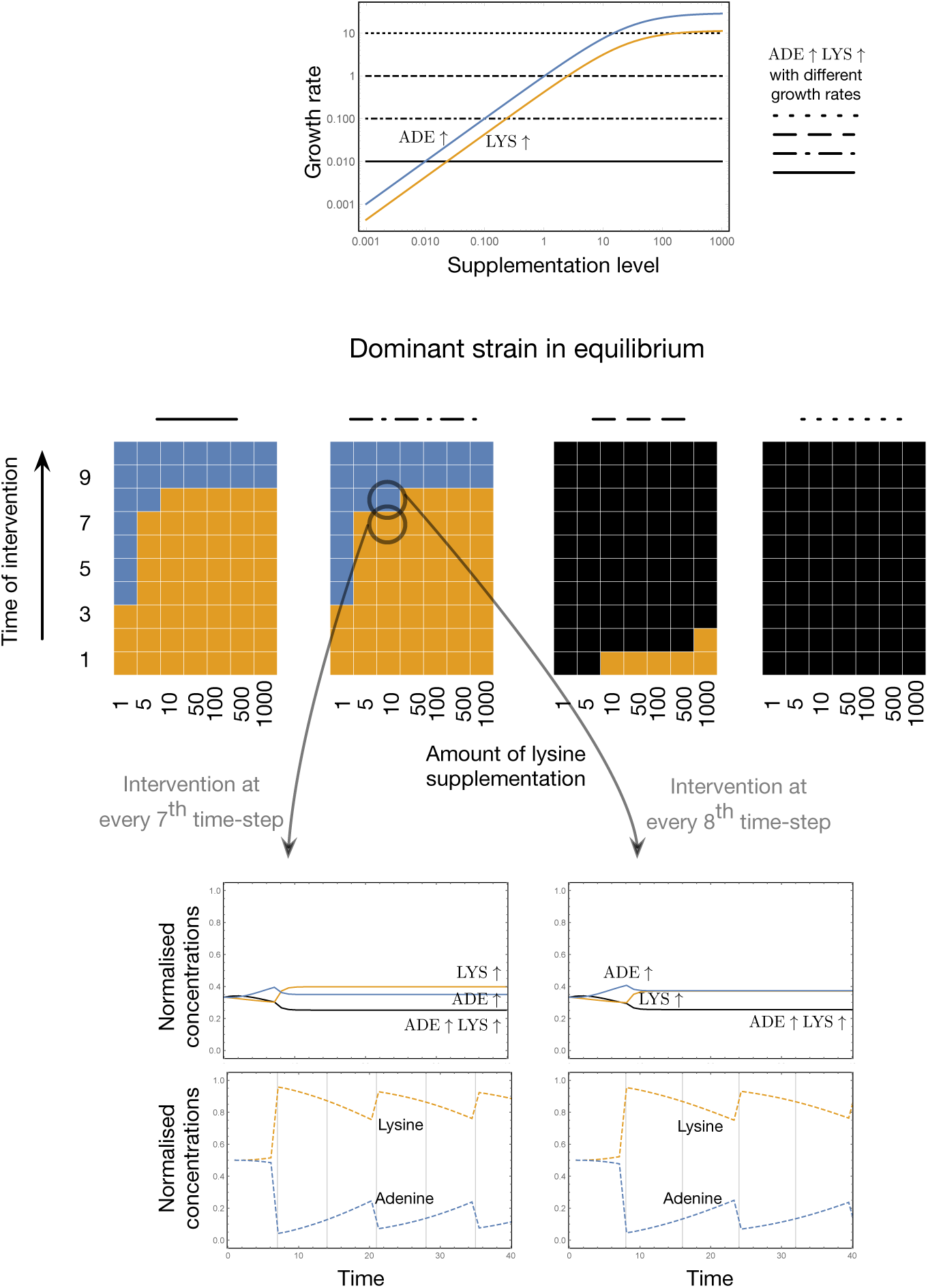
Ultimate overproducers and equilibrium dominance. The top panel displays experimental growth data for the *ADE↑* and *LYS↑* strains. A range of constant growth rates for the *ADE↑LYS↑* strain that fall inside of the aforementioned experimental rates were selected, shown with the varying black lines. Given these growth rates we can estimate the equilibrium of the system. The colour denotes the dominant strain at equilibrium in each simulation, with yellow representing *LYS↑* and blue 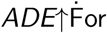 increasing growth rates of the ultimate overproducer, we see that more lysine is required to maintain one of the mutualists as the dominant strain. Furthermore, the exact time of intervention *k* (and then every *k*^th^ step) matters. The low fitness strain *ADE↑LYS↑* never becomes dominant, but relative dominance of the two mutualists can change (left two panels and the explicit dynamics shown in the insets). For reduced supplementation, the intervention must occur earlier if there is to be a cha1n1ge in dominance. The same is true for higher fitness of *ADE↑LYS↑* although in this case, intervention times would hvae to occur extremely early on.

The theoretical model shows that we can supplement the culture with one of the metabolites. Since the *ADE↑* strain has the higher growth rate, we support it by providing lysine in the culture (environment). As the amount of supplementation is reduced, it must be provided earlier so as to maintain *ADE↑* as the dominant strain. The later intervention via supplementation occurs, the more likely that predictions of the growth rate for unsupplemented populations hold true. The same holds for scenarios in which the ultimate overproducer dominates. We need to provide a high amount of supplement early to offset the fitness benefit of the ultimate overproducer Fig. 6 (dominant strain panel 3). If supplementation occurs after the carrying capacity has already been reached, then it has no effect.

## Discussion & Conclusion

Cooperation is instrumental at all levels of life and on all timescales. On a scale of ecosystems, the biosphere itself can be viewed as interacting networks of varying components. Although important, the behaviour of these complex interactions is difficult to study and nearly impossible to predict. Even simple interactions between a small number of cooperators form complex networks, but when these networks are placed in fluctuating environments their complexity surges. Synthetic biological systems apply engineering principles to an organism to promote precise control and predictability over natural behaviour. To this end, we leverage the simplicity of the engineered yeast system and the corresponding mathematical model. Our work examines both the effects of how initial strain ratios and changes in the environment alter the dynamics of strain concentrations and influence mutualism stability. Under stable environmental conditions, as previously shown, the initial strain ratio of the system is very stable[15]. Stability under changing environmental conditions is an entirely different matter. Environmental changes have the potential to influence or disrupt mutualistic interactions [13]. Here we dissect these influences further with the aim of discerning how mutualistic interactions may be maintained. Beyond interactions between biotic components, we have integrated abiotic factors, such as the amount and timing of environmental change in our system.

The final equilibrium of the mutualistic system depends more on environmental supplementation shocks than on the starting ratio of the mutualist partners. As initial supplementation of adenine increases, favouring the *LYS↑* strain, the final ratio shifts in both the experimental system and mathematical model (Fig. 3). Confident in the exploratory power of our mathematical model, we determined the environmental conditions that promote mutualism and provide what we refer to as the Goldilocks zone. The modeled supplementation regimes are common in microbial culturing, but they also have clear ecological parallels. The non-supplemented culture represents a niche baseline. Initial supplementation is akin to an isolated event like the sinking of a whale carcass to the floor of the ocean, resulting in extreme point-source enrichment [28]. Intermittent supplementation represents seasonal or periodic change, such as temperature fluctuations or the regular introduction of a nutrient to the gut of a host animal. Understanding the effect of fluctuating resource availability is extremely important regarding invasiveness [29]. A temporary reduction in competition due to nutrient excess can make mutualisms vulnerable to invasion [30]. Finally, continuous supplementation is a permanent change to an environment, as in a permanent temperature change or evaporation of a large body of water. In general, we identified a gradually shrinking Goldilocks zone as one progresses from initial, to intermittent and then to continuous supplementation regimes (Fig. 4). With environmental destabilisation becoming more and more concrete, the edges of the mutualistic system encroach upon even minimal supplementation levels. Stochastic effects may dominate these regimes in which mutualism can then collapse due to chance events. Tracking the size of the zone is thus a good measure of the resilience of a mutualistic system under changing environments.

The mode of intermittent supplementation and in particular, its timing, offers some respite. These yeast strains have asymmetrical starvation tolerance and do not release their overproduced metabolite until near death [15]. Intermittent supplementation could thus represent death-induced periodic release of nutrients. It is important to note that the frequency of disturbance in complex communities can severely affect assemblages of microbes [31]. We explored this by starting intermittent supplementation at different times and by varying supplementation frequency. SI. highlights the importance of supplementation parameters as well as the starting ratio of the two strains. We observe that when environmental changes are minute, the initial ratios matter somewhat, but if the change in the environment is drastic, then the timing of the change is crucial.

We include the analysis of a hypothetical ultimate overproducer *ADE↑LYS↑* that allows use to explore both the cost of mutualism generally and to tease part the influence of the timing and magnitude of environmental disturbances. Although even modest costs can be rapidly selected against, it is important to recognise that mutualism need not be expensive from a cellular point of view. As such, contributions to the common good may persist as what could be described as genetic anomalies. These individuals are not cheaters, as they support the existing mutualistic interaction, but could still quickly become dominant members of an ecosystem if the cost of contribution is lower than the benefits gained from not relying on other members of the ecosystem (Fig. 6). Moreover, even for a high cost the *ADE↑LYS↑* strain can persist in a stable equilibrium with the other strains, in part because they are not dependent upon cycles in abiotic factors.

The disruptive influence of this overproducing strain can be mitigated. Intervention both early and with considerable supplementation permits continuation of the mutualistic interaction between *ADE↑* and 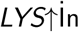 principle, targeted intervention in collapsing ecological niches that depend on mutualism could save these relationships, or at least forestall their collapse. Further experimental tests along these lines will provide insight into the role of abiotic components in the resilience of mutualistic systems.

Understanding environmental enrichment (or degradation) is imperative as we face climate change. The promise of bioengineering together with cooperation is enormous [32]. Beside offering insight into the nature of interactions resulting in community stability [33], engineered systems have implications for designing complete ecosystems that affect the biosphere in general [34]. With continuing eutrophication of oceans, essential symbiosis may break down into parasitism [6]. Use of pesticides and synthetic nitrogenous fertilisers can disrupt the natural nitrogen fixation process – mutualism between leguminous plants and rhizobacteria [4]. Thus, enriched environments could pose a threat to long evolved mutualism which might lead to further catastrophic events [35]. Since mutualisms bind organisms together to enhance their survival, they come at the cost of binding the fates of all involved species together [36]. Changing the environment, inadvertently, or purposefully via anthropogenic activity and climate change can be costly not just to one set of species, but to the ecosystem at large. Beyond conservation biology, mutualistic interactions also lie at the heart of translational biology, affecting applied human health and biotechnology. For example, normal functioning of the human microbiome is decisive for our well-being, and various pathologies emerge from disruption of that microbiome [37].

Microbes often occur as consortia rather than as individual species. Communities of bacteria are usually the pioneers in harsh environments, from newly formed volcanic rock to hyper-arid deserts [38, 39, 40]. Some members of the consortia survive by feeding off inorganic matter, but most of the community survives in a cross-feeding network [41]. In such cases, it would be useful to know how the communities function, and especially their fates when the environment has changed *due to their presence*. While the diversity of individuals in a consortium does not always guarantee robustness, engineered microbial consortia might prove to be more robust in harsh environments [32]. Learning from natural, well-established communities, we can better design interactions between microbes, making them resilient against environmental disturbances, and thus useful tools in biotechnology. When developing techniques to tackle problems such as wastewater treatment, biofuel generation, or oil spill cleanups, symbiotic communities of microorganisms potentially offer a more efficient pathway to the breakdown of complex substrates [42].

Understanding mutualism can therefore help address questions about the origins, spread, and diversification of life in inhospitable environments. Synthetic microbiology will help us take these mutualisms to the next level of rich environments. Competition between multiple, synthetically constructed, cross-feeding systems with different levels of interdependence would help extract information about successful ensembles of interactions eventually leading us to the evolution of successful hypercycles – as envisioned by Eigen and Schuster[20]. Our next steps in synthetic biology will be in this direction.

## Acknowledgments

The authors thank Nicholas Luscombe and David Rogers for a careful reading of the manuscript and providing helpful comments and Steven Aird for editing the manuscript. J.A.D. and C.S.G. acknowledge the generous funding from Okinawa Institute of Science & Technology and from the Max Planck Society respectively. The authors would like to thank Professor W. Shou for providing the yeast system used in this work.

## Author contributions

J.A.D. and C.S.G. conceived the project, analsysed the results and wrote the paper. C.S.G. developed the model. J.A.D. performed the yeast experiments.

## SUPPORTING INFORMATION

### Methods and Materials

#### Saccharomyces cerevisiae Strains

The synthetic system is composed of two metabolite-overproducing strains previously developed in the w303 background, provided by Professor W. Shou [15]. The first, WS950, is an adenine-overproducing strain denominated *ADE↑* throughout this manuscript. It has the genotype MATa ste3::kanMX4 lys2Δ0 ade4::ADE4(PUR6) ADHp-DsRed.T4. The second, WS954, is a lysine-overproducing strain called *LYS↑* here. This strain has the genotype MATa ste3::kanMX4 ade8Δ0 lys21::LYS21(fbr) ADHp-venus-YFP.

#### Culturing & Plate Counts

Synthetic complete (SC) media was made from dextrose, FORMEDIUM yeast nitrogen base and an appropriate synthetic complete dropout supplement (Kaiser). Amino acid free minimal media media (SC-aa) was made as above without synthetic complete dropout supplement. Yeast-extract, peptone, dextrose (YPD) media was made using chemicals from BD Diagnostics or Sigma. All cultures were grown at 30 ^°^C in an orbital shaker at 200RPM.

Experimental cultures were generated by selecting individual colonies from a streak plate and growing each colony in 5 mL SC medium for 48 hours. These cultures were pelleted, washed twice with SC-aa and resuspended in 5 mL of SC-aa. They were then grown for an additional 24 hours. These cultures were pelleted, washed and resuspended as above and then were diluted 1 in 20 with SC-aa. Each experiment was performed in triplicate starting from a single individual colony. Single-strain growth tests were performed using 5 mL aliquots of the 1 in 20 dilution and supplementing to a final concentration with either 0, 0.1, 1, 10, or 100 µg/mL of adenine for the *LYS↑* strain or lysine for the *ADE↑* strain. The cultures were sampled at 0 and 24 hours. Experimental cultures were generated by mixing individual strain dilutions at volume ratios indicated in each experiment to a final volume of 5 mL. The, 0, 0.1, 1, 10 or 100 µg/mL adenine-supplemented cultures peaked in the number of colony forming units, suggesting growth saturation, at 144, 120, 72, and 48 hours respectively. In each case, an equilibrium was reached before saturation. We determined colony forming units via plate counts on YPD solid media of culture dilutions at time points indicated. Strain ratios were determined by replicate plating these YPD colonies onto solid media that permitted growth of only one strain, either SC without lysine or SC without adenine, and counting the respective colonies.

**Estimating growth parameters.** We now parametrize the Michaelis constants (*k*_*c*_*A* and *k*_*c*_*L*) and the growth parameters (*r*_1_ and *r*_2_) using experimental data. Single-strain growth rates for *ADE↑* and *LYS↑* were determined on growth media supplemented with varying concentrations of the appropriate required metabolite. The two strains were grown in synthetic complete media either unsupplemented or supplemented with 0.1, 1, 10, or 100µg/mL of the corresponding nutrient. Growth after 24 hours was calculated as above. The growth curves were used to parameterise model growth components.

#### Data Analysis

Colony counts were collated and summary statistics generated using R [43]. A RMarkdown file containing raw data, summary statistics, all code and preliminary plots is available on GitHub at https://github.com/tecoevo/syntheticmutualism and will be included as a supplementary file in the final submission. All figures were generated using Mathematica[44] for consistency.

#### Model extension for supplementation regimes

For initial supplementation we do not need to modify the model from what its form in Eqs. (2) and (4). For different values of initial supplementation, we change the initial conditions for the amounts of nutrients. For *continuous supplementation* we modify the Eqs. (4) to,

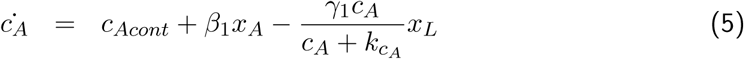

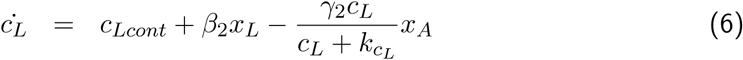

where the values of *c*_*Acont*_ and *c*_*Lcont*_ determine the amount of nutrient continuously added to the culture. For *intermittent supplementation* we make use of a hybrid dynamic system [23]. We start with the same initial conditions as for other supplementation regimes, where the initial condition of the nutrient matches the amount used for supplementation. Dynamics proceed as per equations in the main text. In total, they run for the same amount of time as the initial and continuous supplementation experiments before assuming equilibrium, but the time is split into many small cycles. Each cycle runs for a short period. Thus we have *cycle length×number of cycles* = *total time until equilibrium*. At the end of each cycle, we add the predetermined amount of essential metabolites and then allow the next cycle to continue. In the absence of external supplementation, the system requires a certain time to equilibrate. If supplementation continues to disrupt this process by disturbing transient dynamics, then the eventual outcome when evaluated at the same time point as in other supplementation regimes can be drastically affected (Fig. 5 and Appendix Fig. 7).

**Figure 7:**
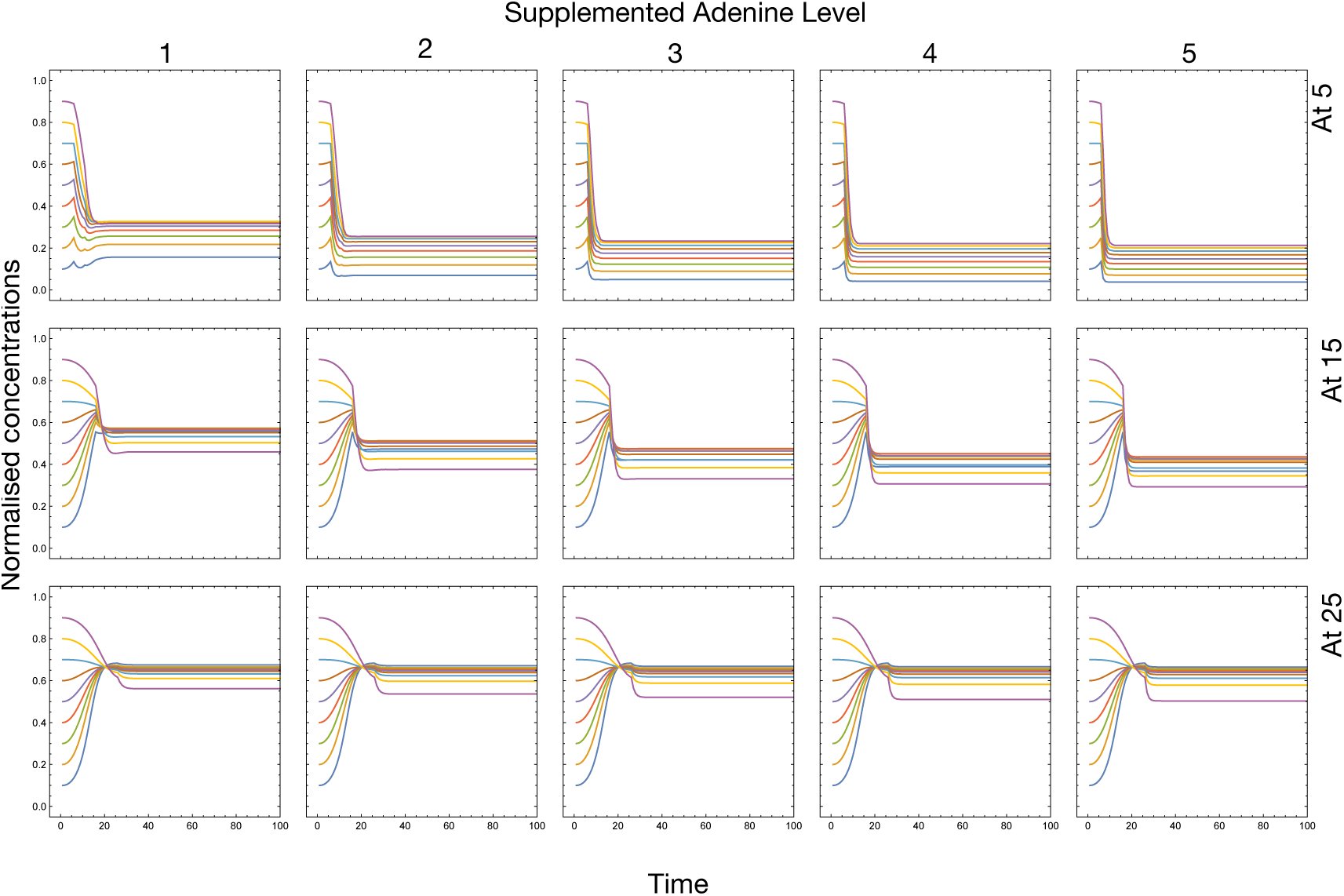
As shown in Fig. 5, the frequency and timing of intermittent supplementation changes the equilibrium of the system, but the initial conditions matter as well. In this figure we highlight this dependence. Starting the supplementation regime at time point 5 and then supplementing at every 5^*th*^ time point we get the first row for different levels of supplementation. For the second and third row, the first dose (and subsequent) of supplementation occurs at 15 and 25 time-points. For high supplementation, the number of cycles are immaterial. A single dose of supplementation is enough to shift the equilibrium; however, the exact timing of this dose is crucial. The time at which the first disturbance occurs affects the role of initial conditions. If supplementation begins early (at 5), then the order of initial conditions is not affected as opposed to cases in which supplementation is started later (at 15 and at 25).

